# stLFRsv: a germline SV analysis pipeline using co-barcoded reads

**DOI:** 10.1101/2020.06.15.141721

**Authors:** Junfu Guo, Chang Shi, Xi Chen, Ou Wang, Ping Liu, Huanming Yang, Xun Xu, Wenwei Zhang, Hongmei Zhu

## Abstract

Co-barcoded reads originated from long DNA fragment (mean length larger than 50Kbp) with barcodes, maintain both single base level accuracy and long range genomic information. We propose a pipeline stLFRsv to detect structure variation using co-barcoded reads. stLFRsv identifies abnormally large gaps between co-barcoded reads to detect potential breakpoints and reconstruct complex structure variations. The barcodes enabled co-barcoded reads phasing increases the signal to noise ratio and barcode sharing profiles are used to filter out false positives. We integrate the short reads SV caller smoove for smaller variations with stLFRsv. The integrated pipeline was evaluated on the well characterized genome HG002/NA24385 and obtained precision and recall rate of 74.2% and 22.3% for deletion on the whole genome. stLFR found some large variations not included in the benchmark set and verified by means of long reads or assembly. Our work indicates that co-barcoded reads technology has the potential to improve genome completeness.

## Introduction

Structure variations (SVs) which represent genome variations larger than 50bp, consisting of deletions, insertions, inversions, tandem duplications and translocations. They contribute more difference than SNP and small indels between genomes (Pang 2010). Some of them are pathogenic variations and associated with specific diseases (Jongmans 2006, Singleton 2003, Rovelet 2006). Despite the importance of SVs, profiling them has been challenging.

For the recent 20 years, many technologies have made SV annotation move forward and made available a well-characterized human genome to facilitate the developing of SV identification tools (Justin 2019). Among these technologies sequencing is a main category, which includes short read, long read and co-barcoded read sequencing. Each sequence technique, with its unique advantages and disadvantages, contributes to uncovering the SV profiles among population.

Short reads are accurate at base level and cost effective. Their uniform depth and insert-size are successfully used to identify deletions and insertions, or copy number variations (Layer 2014, Talevich 2014). Deletions are easier to detect than insertions. More complex ones seldom present in the results because their breakpoints are usually in close proximity to regions lack of unique short read alignment.

Long reads or single molecule sequencing reads are usually with length N50 above 10Kbp. They catch breakpoints easier and may go across repetitive regions nearby (Jain 2018). They prone to have insertion and deletion errors and the base level accuracy is comparatively low, which leads to the low accuracy of small variation (below 200bp) detection (Wang 2019). Single molecule circular consensus sequencing protocol produces highly accurate and averagely above 10Kbp reads (Wenger 2019). However, it is not applicable to large scale projects yet because of the output and cost limit.

Co-barcoded reads are the product of novel protocols for library construction and NGS sequencing technology (Wang 2019, Zheng 2016, Zhang 2017). This strategy enables all the short reads originating from each long DNA molecule to share a common barcode. Thus they gain long range genome information while maintaining base level accuracy. The nanograms of input DNA make it feasible to many applications. The inferred averaged DNA fragment length for co-barcoded reads usually reaches 50Kbp, which makes it possible to sequence across even larger repetitive sequences nearby SV breakpoints.

Analysis pipelines that detect SVs utilizing co-barcoded reads fall into three categories in respect of ways they use barcode information. The first category identifies novel adjacency by detecting abnormal numbers of common barcodes shared between two genome loci or bins (Marks 2019, Li 2018, Spies 2017). The second one tests the distribution of sequenced short segments on large DNA molecules (Marks 2019, Rebecca 2017). The third uses barcode numbers to extract data for local assembly (Zhou 2019, Meleshko 2019).

Here we present stLFRsv, a co-barcoded reads SV analysis pipeline, which falls into the first category and integrates short reads SV detector smoove (https://github.com/brentp/smoove).

## Results

The HG002 cell line sample was processed according to single tube long fragment read (stLFR) protocol (Wang 2019) and sequenced to 100x coverage. The average number of read pairs per barcode is 51. The inferred mean fragment length is 83Kbp. And the inferred mean number of fragments per barcode is 1.15. The distributions of read pair numbers, weighted fragment lengths and fragment number per barcode are illustrated at supplementary Fig. 1. We down sampled the data to 50X and 30X and called variants separately to provide guidance for applications. stLFRsv was assessed on HG002 genome against four SV callers: Long Ranger, NAIBR, smoove and GROC_SVs (Marks 2019, Rebecca 2018, Spies 2017). And the results from co-barcoded reads were compared with structure variations from 100X Nanopore long reads. The commands used to run the following pipelines are shown by Supplementary Table 1.

### Structure variation from co-barcoded reads

The workflow of structure variation detecting is illustrated in Fig.1A. Co-barcoded reads were aligned to hs37d5 by BWA-MEM2 (Li 2013). Phasing was made by Hapcut2 after SNPs were called using GATK (Edge 2017, McKenna 2010). The GIAB v0.6.0 structure variation set includes 7172 insertions and 5336 deletions. We used Truvari to align pipeline calls to GIAB call set (https://github.com/spiralgenetics/truvari). For Long Ranger the alignment was done by Lariat. For other software, the alignment results by BWA-MEM2 were used.

**Fig. 1.**
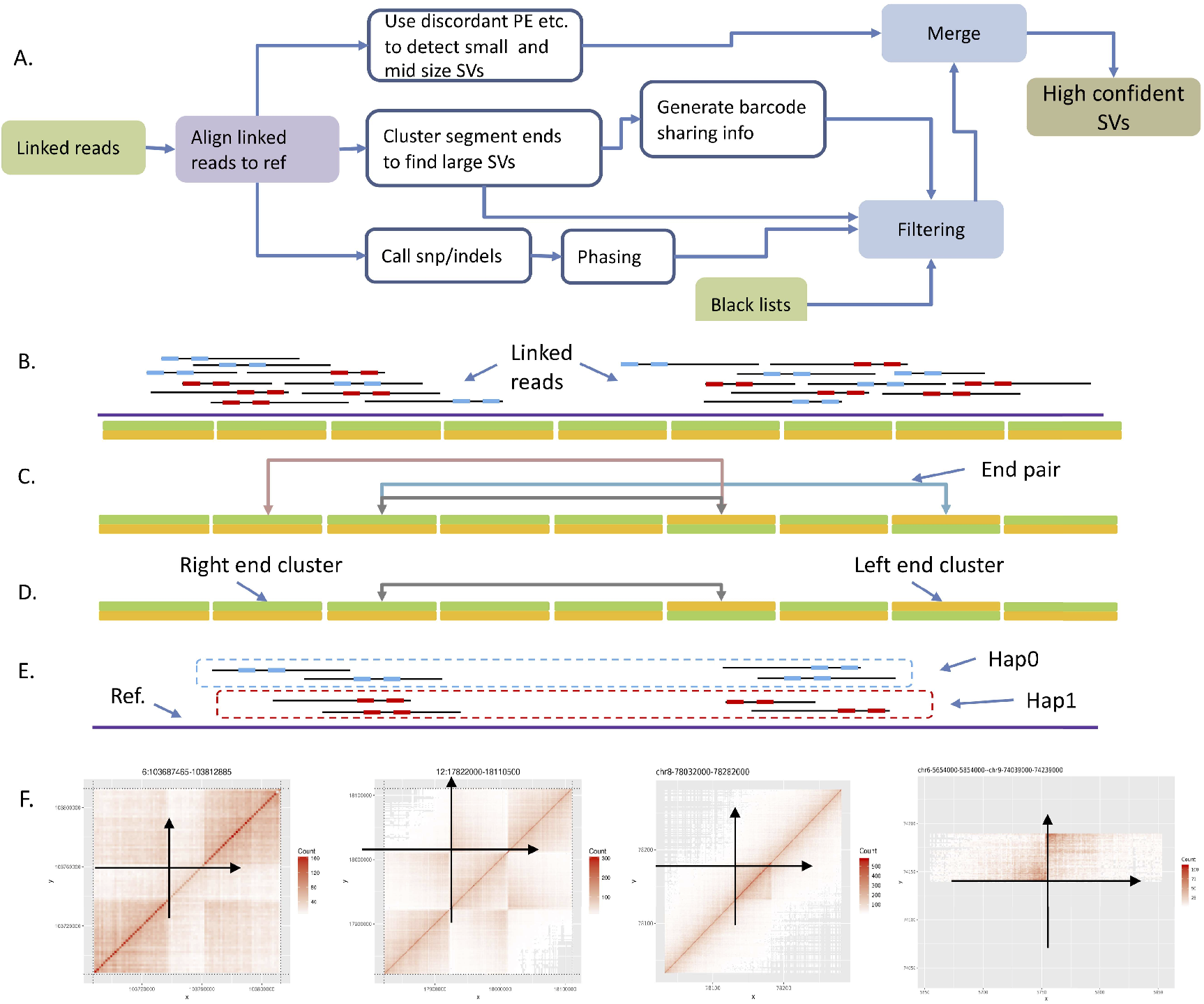
Workflow and algorithm. **a**, Structure variation detecting workflow. **b**, Cluster segment ends by bins: left end cluster and right end cluster. **c**, Pair up ends by shared barcodes. **d**, Pare down candidates by removing those nearby. **e**, Split by haplotypes through phasing barcodes on phasing blocks. **f**, Use barcode sharing heatmap pattern as a filter and anchor the variation on the genome.

78 large deletions were identified by stLFRsv. 32 of them were validated by GIAB call set. Among the 46 unmatched deletions, 14 of them overlap with the GIAB deletion records but were failed by Truvari. 28 of them overlap with the GIAB deletions with markers other than “PASS” (Supplementary Table 2). One of them is located at Chr3:195,196,500 – Chr3:195,213,000 (Fig. 2C, 2F). In this area the Nanopore assembly indicated an approximately 30Kbp alternative sequence to the reference genome. Four deletions do not overlap any GIAB record. One of them is located at Chr16:18,828,000 - 18,840,000 and confirmed by Nanopore long reads (Fig. 2A, 2E). Another is located at Chr19:21,823,500 - 21,835,500 and indicated as a heterozygous variation by long reads (Fig. 2G).

**Fig. 2.**
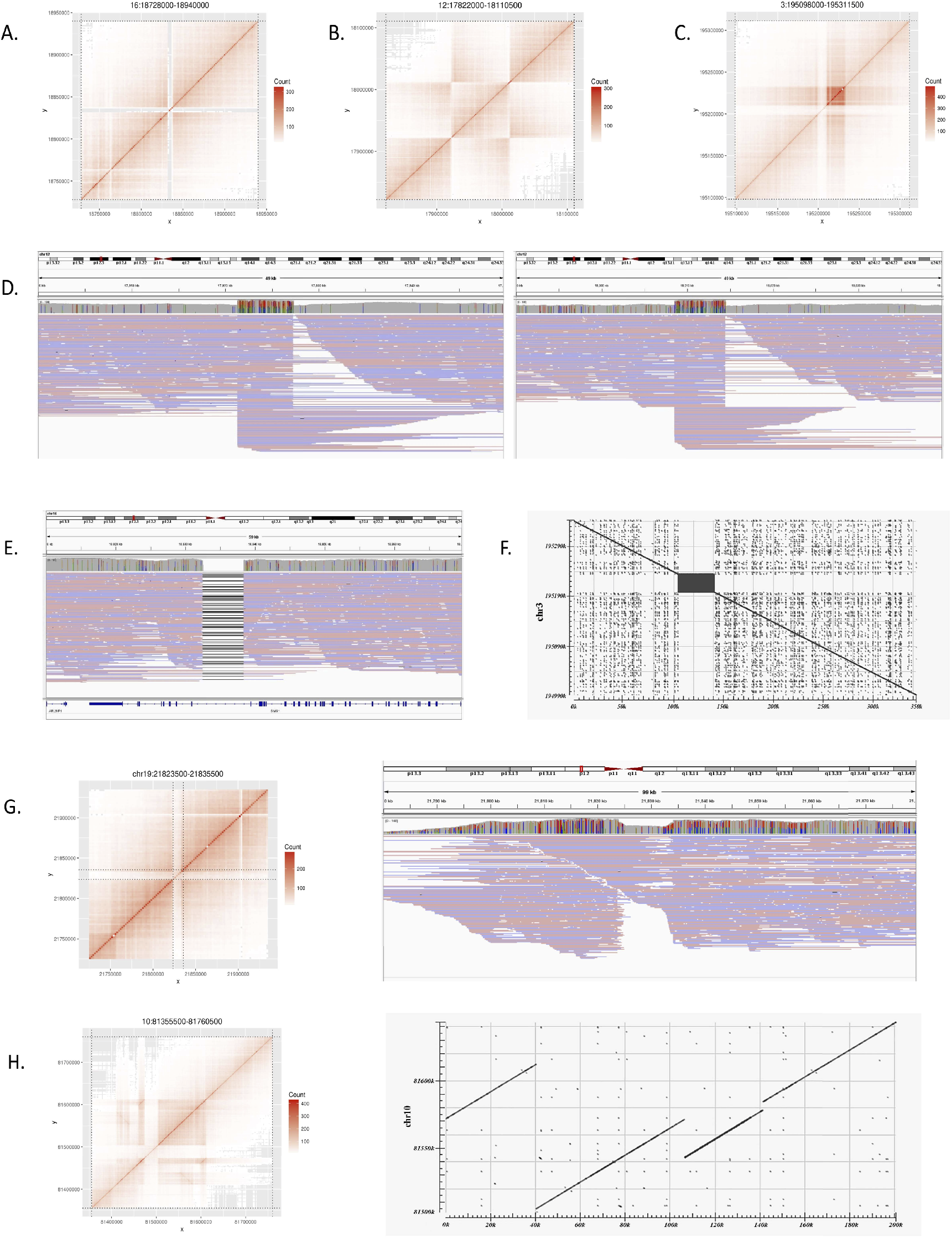
Large variations not match the GIAB benchmark. **a**, Heatmap for a deletion on Chr6. **b**, Heatmap for an inversion on Chr12. **c**, Heatmap for a deletion on Chr3. **d**, Long reads alignment supports the inversion in b. **e**, Long reads alignment supports the deletion in a. **f**, Assembly alignment to reference by Blast for the deletion in c. **g**, Heatmap for a deletion on Chr19 and long reads alignment. **h**, Heatmap for a duplication on Chr10 and assembly alignment.

Eight GIAB deletions larger than 10K were not detected by stLFRsv whose heatmaps are in Supplementary Fig. 2. Two of them are in the N-regions of the reference. Five of them were undetected because of confusing signals or lack of signals across the deleted area. One of them is actually an inversion in which nested a smaller inversion.

Other than deletions, stLFRsv found 12 inversions, duplications and translocations (Supplementary Table 3). All of them are shared by multiple genomes which indicates problematic reference regions or repeat sequences on reference. For example, one inversion was found at Chr12:17,922,000 – Chr12:18,013,500 (Fig. 2B, 2D). It was classified to be a homozygous variant and confirmed by Nanopore reads. One duplication was identified at Chr10: 81,505,500-81,610,500, which was confirmed by long reads assembly (Fig. 2H).

When merging deletions from stLFRsv and smoove, the size cutoff was set to 10Kbp by stLFRsv based on the data profiles. The deletion evaluation results are shown in Table 1. The down sampled results are in Supplementary Table 4. Since few insertions were found by any of the four callers, we didn’t perform insertion results evaluation.

**Table 1.**
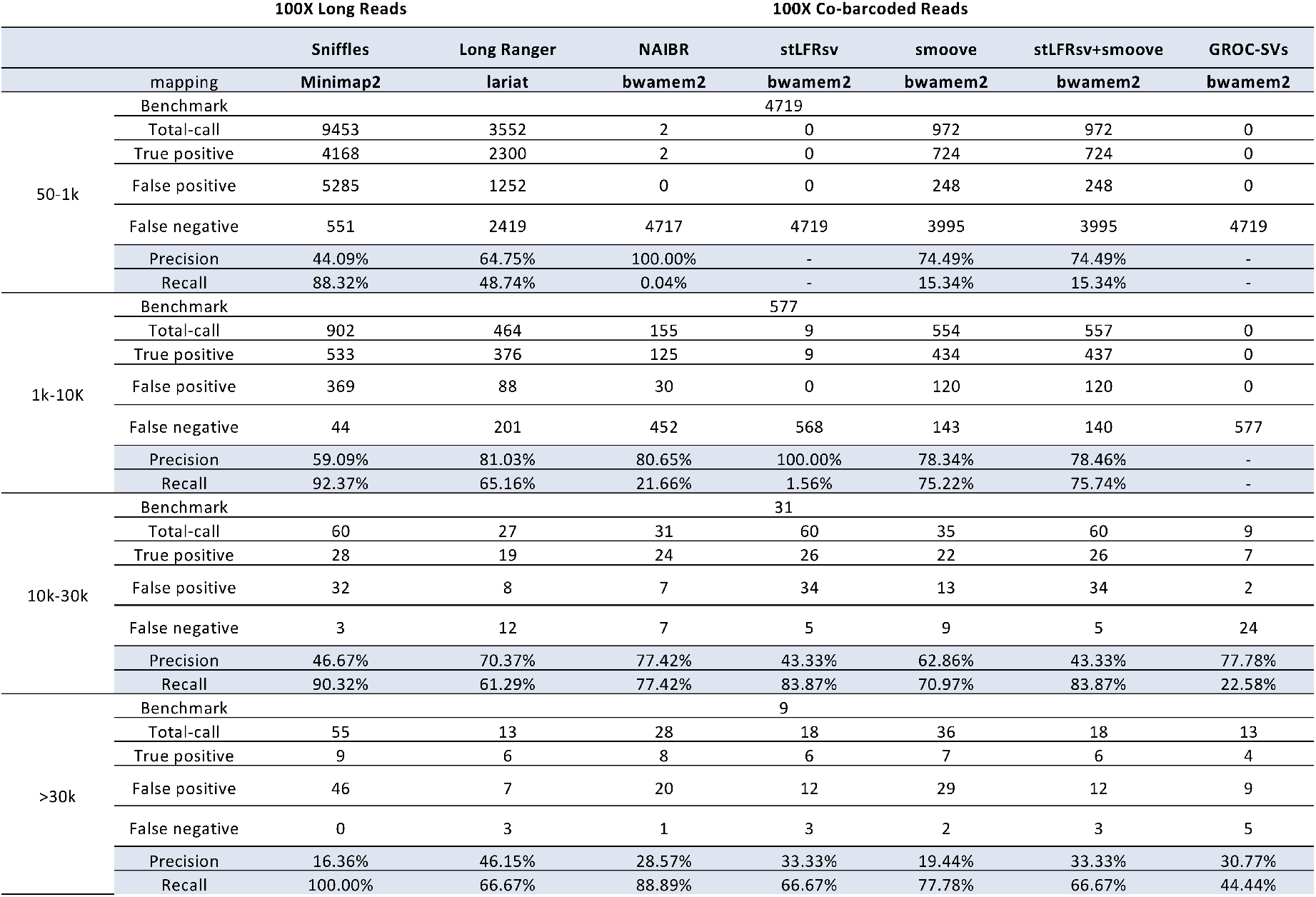
Deletion evaluation on whole genome against GIAB HG002 benchmark.

### Comparing with Nanopore long reads

We got 100X Nanopore long reads of HG002 from Oxford Nanopore Technologies. The distribution of read length and percent identity is resented at Supplementary Fig. 1. The alignment was done by Minimap2 with default parameters (Li 2018). SVs were detected by Sniffles with default parameters while rising support read number can reduce both false positives and true positives (Sedlazeck 2018). The deletion evaluation is also listed in Table 1. And the insertion evaluation is shown in Supplementary Table 5. We assembled long reads by NECAT for variation validation purpose (Chen 2020).

## Methods

Large structure variants leave large gaps in long fragments as viewed from co-barcoded reads alignment (Fig.3). The distribution of read pairs on long fragments is approximately random and the gap sizes between read pairs vary in a wide range. The large gaps appear on long fragments by chance. However large structure variants are likely to lead to large gap aggregation. So stLFRsv detects large gaps in fragments to find large variants. On the other hand, smoove is a pipeline that uses LUMPY as its core to catch pair end discordance and other short reads signals to identify variants (Layer 2014). We use smoove to find small and mid-sized variants. There may be overlap between the two variant sets and we merge the variants before outputting the results (Fig. 1A). The detecting process by stLFRsv is described in the following.

**Fig. 3.**
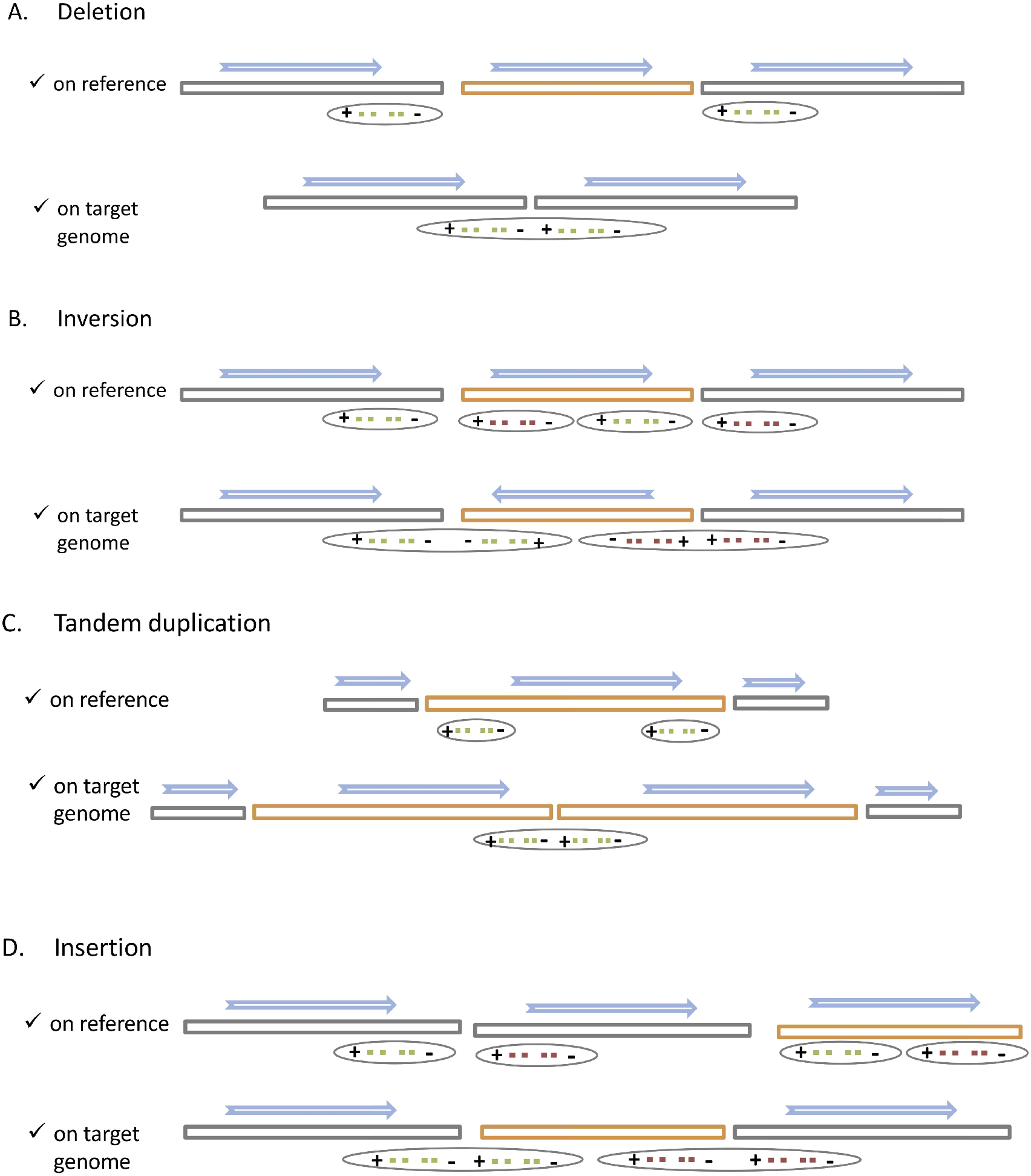
Large structure variations induced large gaps break long fragments into sub-fragments. **a**, Deletions. **b**, Inversion. **c**, Tandem duplication. **d**, Insertion.

### Cluster segment-ends

We calculate an empirical gap size distribution and select a size value *G* as cutoff so that the probability of gap sizes smaller than *G* is 98%. When we break a long fragment at gaps larger than *G*, we get at least two sub-fragments. We define left and right ends as the left and right ends of a sub-fragment. Each end has its position on the reference. We divide the reference sequences into consecutive bins. Each bin has a size of *B* Kbp based on the data profile, which corresponds to a left and a right end cluster. All end clusters with at least one end are kept for the next step (Fig. 1B).

### Pair up ends

If two end clusters have barcodes in common, they form an end pair (Fig. 1C). These end pairs are potential novo adjacencies. We then make an empirical distribution for common barcode numbers of all end pairs and also model common barcode numbers by a binomial distribution with parameters estimated from the same data. With the two distributions and a user set percent value, we obtain a common barcode number threshold for screening end pairs as novo adjacency candidates. There are four types of end pairs according to the types of the two end clusters. If the potential novo adjacency does not involve an orientation change, the end pair is a right-left or left-right. Otherwise it is either a left-left or a right-right type (Fig.3). If an end cluster is in pair with multiple clusters and one of the pairs is very likely the other end of its sub-fragment, we unpair them.

### Pare down candidates

Since the DNA molecules are partially sequenced, sub-fragment ends do not gather densely around a novo adjacency. They may spread in several bins and give arise to multiple end pairs.

To reduce redundancy, for each end pair, we compare its common barcode number with those of the pairs in the same type whose ends locate no more than four bins away and only keep the one with the highest (Fig. 1D).

### Split by haplotypes

About 60% reads are haplotype solved, which means those reads along with their barcodes are placed onto one of the haplotypes of each phasing block (Fig. 1E). We decide whether an end cluster is homozygous or heterozygous using the Fisher’s exact test on the numbers of barcodes phased to each of the two haplotypes with the expected value being half of the haplotype-solved barcodes. If two ends of a pair on the same phasing block, the pair’s zygosity is decided by another Fisher’s exact test on the haplotype-solved sharing barcodes between the two ends with the expected value being half the total. If two ends of a pair are on different phasing blocks, its common barcodes should support mainly one or two of the four possible haplotype combinations. The zygosity of an end pair and whether the zygosities of the two ends match are important information for their confidence level.

### Filter

Noisy signals often result in false novo adjacencies. Here are some noise filters to mitigate the problem. Firstly we check read mapping quality in the two ends of each pair and screen out the pairs when low quality ratio is above a cutoff. Secondly we filter out candidate pairs in the problematic regions of the reference. These regions are formed based on the reference profile and usually involve repeat sequences, mis-assembled areas and gaps.

The third filter uses the common barcode heatmap around each novo adjacency region (Fig. 1F). A novo adjacency gives rise to the number of common barcodes. This rise shows specific patterns in the regions in close proximity to the novo adjacency on the heatmap. Since this is not a graphic detector, we digitalize the heatmap in order to catch the patterns. If the two directions intersect at the breakpoints on the heatmap, it forms four regions. For a deletion, insertion or duplication, there is only one region showing typical adjacency barcode sharing. For an inversion, there are two regions with symmetric sharing. We collect bin-to-bin sharing numbers in each region and do the Wilcoxon signed-rank test to verify the expected patterns between four regions.

### Anchor breakpoints

If a novo adjacency is formed by a pair of ends which are far away from each other on the reference, we would like to know whether this rearrangement results in a short interrupt or a long-range structural variation. Due to the limited DNA fragment lengths, the numbers of shared barcode fade out as bins moving away from the novo adjacency. When end pairs are placed on the target genome, they all present as a right and left end. If we check the common barcode numbers between the bin holding the right end and the bin holding the left end and bins following the left, the common barcode numbers should show a gradual decline. We calculate the difference between observed numbers and the expected numbers according to the distribution described above. The process is similar for the left end. For each end pair we have two lists of differences and test them by a Wilcoxon signed-rank test to detect a sudden loss of barcode sharing in one of the directions. If there’s evidence supporting a sudden loss, we infer it’s a short sequence from this direction inserted to the other direction. This estimation is more accurate for haplotype-solved novo adjacencies.

### Merge

We extract variations below a cutoff size from smoove results and those above this cutoff from stLFRsv results and combine them by merging those with significant overlap (at least 70% overlap with respect to the longer one) to form the final output.

## Discussion

We present stLFRsv a co-barcoded reads structure variation detector which finds large variations with much less false positives than detectors not using long range information. When combined with standard short reads variation caller, it can exploit the co-barcoded reads to uncover the whole spectrum of genome polymorphism.

More large variations other than deletions and insertions are needed to assess the ability of variation detection by co-barcoded reads. We are analyzing co-barcoded reads for clinical samples to find balanced translocation. This technique is likely to give more precise description of such variations than current clinical practices. And another clinical application is to associate a genetic defect with nearby alleles using co-barcoded reads to infer whether the infant gets the defect through pre-natal cell free DNA sequencing by detecting its associated nearby alleles. This one makes use of the phasing ability of co-barcoded reads. Another future work is to add a local assembly module to enhance small variation detection (at the range of 50bp~1Kbp).

At the cost a little higher than standard short reads, co-barcoded reads are able to reveal much more useful information of the underlying genomes.

## Availability of data and materials

The structure variation caller stLFRsv is available on GitHub (https://github.com/BGI-biotools/stLFRsv). The script to merge smoove and stLFRsv SVs can be found at https://github.com/BGI-biotools/stLFRsv/tree/master/tools. The co-barcoded reads of HG002 is available at https://db.cngb.org/search/run/CNR0026818/. GIAB benchmark is obtained from ftp://ftp-trace.ncbi.nlm.nih.gov/giab/ftp/data/AshkenazimTrio/analysis/NIST_SVs_Integration_v0.6/.

