## Supplemental Figure 1 for "stLFRsv: a germline SV analysis pipeline using co-barcoded reads"

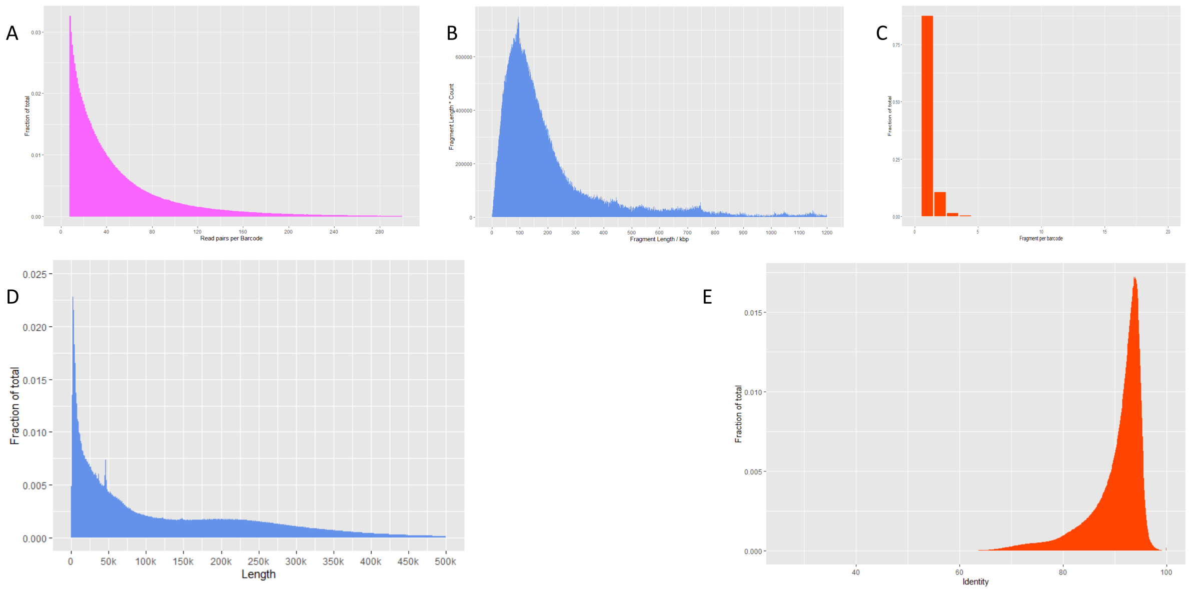


**Supplementary Figure 1. Basic data profile distribution for co-barcoded reads and long reads.**

(a) Read pairs per barcode (b) Weighted fragment length for co-barcoded reads (c) Number of fragments per barcode (d) Long read length (e) Long read percent identity

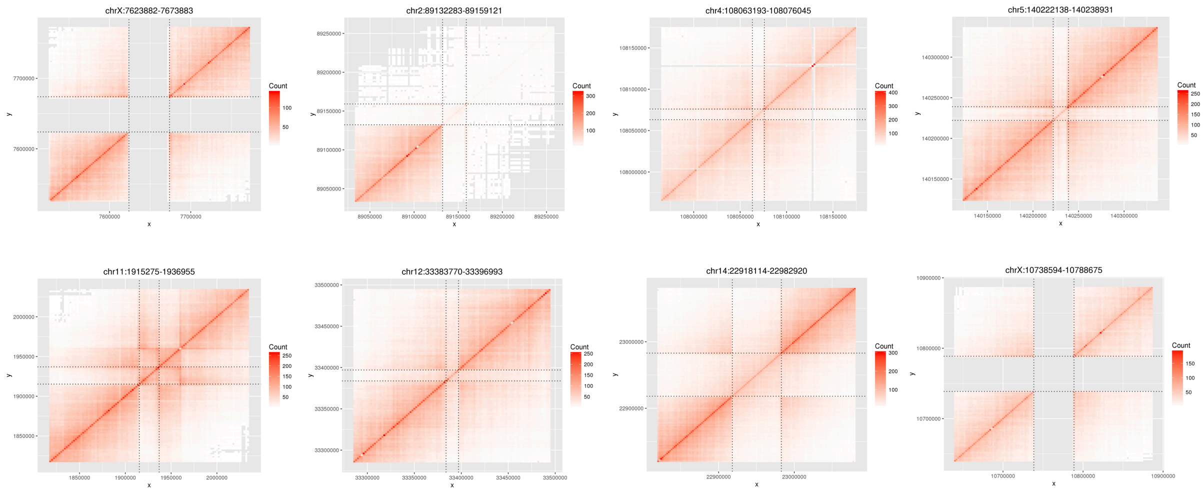


**Supplementary Figure 2. Heatmap for false negative large deletions.**

| | Software | Version | Parameters | Data set |
|---|---|---|---|---|
| short reads | longranger | 2.2.2 | --downsample=130 --vcmode gatk:/.../gatk-package-4.0.7.0-local.jar | 100X co-barcoded reads |
| | NAIBR | - | min_sv=1000 | |
| | grocsv | v0.2.5 | "cluster_settings":{ "cluster_type":"multiprocessing", "processes":10 } | |
| | smoove | v0.2.5 | | |
| | stLFRsv | 2.1 | -size 10000 -gap 10000 -bin 1500 -merge1 4 -merge2 4 -human Y -low2 0.9999 -low1 0.95 | |
| | stLFRsv | 2.1 | -size 10000 -gap 17000 -bin 2000 -merge1 4 -merge2 4 -human Y -low2 0.9995 -low1 0.95 | down_sampled_50x |
| | stLFRsv | 2.1 | -size 10000 -gap 22000 -bin 3000 -merge1 4 -merge2 4 -human Y -low2 0.9995 -low1 0.95 | down_sampled_30x |
| long reads | sniffle | 1.0.11 | --min_support 40 --num_reads_report -1 --genotype --cluster --report-seq | |
| | minimap2 | 2.12-r845-dirty | -ax map-ont | |
| | ngmlr | 0.2.7 | -x ont | |
| | bwa-mem2 | 2.0pre1 | -R '@RG\tID:L0\tLB:COMPLETE\tSM:L0' | |
| | truvari | 1.3.4 | -s=50 --sizemax=200000 | |

### Supplementary Table 1. Parameter setting for the used pipelines

| Chr | Pos | SV type | Marker | Profile | Genotype | Overlap with GIAB records ( overlap length to both SV length ratio > 0.5) | Overlap with GIAB records ( overlap length to both SV length ratio > 0.4) |
|---|---|---|---|---|---|---|---|
| 1 | 72765000 | <DEL> | COMMON | END=72813000;SVTYPE=DEL;SVLEN=-48000 | 0/1 | chr1 72766323 72811839 45516 DEL ClusteredCalls | |
| 1 | 247846500 | <DEL> | PASS | END=247858500;SVTYPE=DEL;SVLEN=-12000 | ./. | chr1 247850455 247856508 6053 DEL PASS | |
| 2 | 1216500 | <DEL> | PASS | END=1228500;SVTYPE=DEL;SVLEN=-12000 | 0/1 | chr2 1218111 1226373 7994 DEL ClusteredCalls | |
| 3 | 129762000 | <DEL> | COMMON | END=129807000;SVTYPE=DEL;SVLEN=-45000 | 1/0 | chr3 129763380 129806745 43365 DEL NoConsensusGT | |
| 3 | 195196500 | <DEL> | PASS | END=195213000;SVTYPE=DEL;SVLEN=-16500 | ./. | | |
| 4 | 9456000 | <DEL> | COMMON | END=9483000;SVTYPE=DEL;SVLEN=-27000 | 0/1 | chr4 9452493 9476819 24326 DEL LongReadHomRef | |
| 4 | 69372000 | <DEL> | COMMON | END=69492000;SVTYPE=DEL;SVLEN=-120000 | 1/0 | chr4 69370648 69487949 117301 DEL LongReadHomRef | |
| 4 | 159273000 | <DEL> | PASS | END=159285000;SVTYPE=DEL;SVLEN=-12000 | 0/1 | chr4 159274254 159284292 8380 DEL PASS | |
| 5 | 155475000 | <DEL> | PASS | END=155496000;SVTYPE=DEL;SVLEN=-21000 | ./. | chr5 155474830 155488939 14109 DEL LongReadHomRef | |
| 6 | 31278000 | <DEL> | PASS | END=31290000;SVTYPE=DEL;SVLEN=-12000 | 0/1 | chr6 31279008 31288711 9664 DEL NoConsensusGT | |
| 6 | 86706000 | <DEL> | PASS | END=86718000;SVTYPE=DEL;SVLEN=-12000 | 1/1 | chr6 86708739 86714791 6052 DEL ClusteredCalls | |
| 6 | 129318000 | <DEL> | PASS | END=129330000;SVTYPE=DEL;SVLEN=-12000 | 1/1 | chr6 129319515 129325574 6059 DEL PASS | |
| 7 | 38386500 | <DEL> | PASS | END=38398500;SVTYPE=DEL;SVLEN=-12000 | 1/0 | chr7 38385673 38394814 9141 DEL LongReadHomRef | |
| 7 | 65053500 | <DEL> | COMMON | END=65097000;SVTYPE=DEL;SVLEN=-43500 | 1/1 | | |
| 7 | 97393500 | <DEL> | PASS | END=97404000;SVTYPE=DEL;SVLEN=-10500 | 1/1 | chr7 97395304 97402639 7335 DEL NoConsensusGT | |
| 7 | 101995500 | <DEL> | COMMON | END=102115500;SVTYPE=DEL;SVLEN=-120000 | ./. | | |
| 8 | 1339500 | <DEL> | COMMON | END=1357500;SVTYPE=DEL;SVLEN=-18000 | ./. | chr8 1341179 1354410 13231 DEL NoConsensusGT<br>chr8 1343263 1351536 8273 DEL LongReadHomRef | |
| 8 | 12385500 | <DEL> | PASS | END=12397500;SVTYPE=DEL;SVLEN=-12000 | ./. | | chr8 12388513 12394492 5979 DEL NoConsensusGT |
| 8 | 24970500 | <DEL> | COMMON | END=24991500;SVTYPE=DEL;SVLEN=-21000 | 0/1 | chr8 24972432 24990943 18511 DEL ClusteredCalls | |
| 8 | 73783500 | <DEL> | PASS | END=73795500;SVTYPE=DEL;SVLEN=-12000 | 1/1 | chr8 73787765 73793823 6058 DEL PASS | |
| 8 | 135078000 | <DEL> | PASS | END=135090000;SVTYPE=DEL;SVLEN=-12000 | ./. | chr8 135082913 135089015 6102 DEL PASS | |
| 10 | 6408000 | <DEL> | PASS | END=6421500;SVTYPE=DEL;SVLEN=-13500 | 1/1 | | chr10 6411563 6417629 6066 DEL PASS |
| 10 | 81472500 | <DEL> | COMMON | END=81505500;SVTYPE=DEL;SVLEN=-33000 | ./. | chr10 81474043 81505183 31140 DEL NoConsensusGT | |
| 10 | 111570000 | <DEL> | PASS | END=111582000;SVTYPE=DEL;SVLEN=-12000 | 1/1 | chr10 111572110 111578216 6106 DEL PASS | |
| 11 | 93150000 | <DEL> | PASS | END=93162000;SVTYPE=DEL;SVLEN=-12000 | 1/1 | chr11 93154136 93160197 6061 DEL PASS | |
| 11 | 95166000 | <DEL> | PASS | END=95178000;SVTYPE=DEL;SVLEN=-12000 | 1/1 | chr11 95169369 95175420 6051 DEL PASS | |
| 12 | 8557500 | <DEL> | PASS | END=8592000;SVTYPE=DEL;SVLEN=-34500 | ./. | chr12 8558485 8590846 32361 DEL NoConsensusGT | |
| 12 | 9631500 | <DEL> | COMMON | END=9735000;SVTYPE=DEL;SVLEN=-103500 | ./. | chr12 9632915 9732036 96158 DEL NoConsensusGT | |
| 12 | 11220000 | <DEL> | COMMON | END=11251500;SVTYPE=DEL;SVLEN=-31500 | ./. | chr12 11219200 11250749 5545 DEL NoConsensusGT | |
| 12 | 13543500 | <DEL> | PASS | END=13554000;SVTYPE=DEL;SVLEN=-10500 | 1/1 | chr12 13545544 13551618 6074 DEL PASS | |
| 14 | 73996500 | <DEL> | COMMON | END=74025000;SVTYPE=DEL;SVLEN=-28500 | ./. | chr14 73996082 74023809 27727 DEL NoConsensusGT | |
| 15 | 20580000 | <DEL> | COMMON | END=20590500;SVTYPE=DEL;SVLEN=-10500 | ./. | chr15 20580643 20588797 8154 DEL NoConsensusGT | |
| 15 | 22335000 | <DEL> | PASS | END=22345500;SVTYPE=DEL;SVLEN=-10500 | 1/1 | chr15 22336854 22344093 7239 DEL NoConsensusGT | |
| 15 | 22372500 | <DEL> | PASS | END=22384500;SVTYPE=DEL;SVLEN=-12000 | 0/1 | chr15 22373078 22383652 10574 DEL NoConsensusGT | |
| 15 | 55216500 | <DEL> | PASS | END=55230000;SVTYPE=DEL;SVLEN=-13500 | 1/1 | | chr15 55218215 55224424 6209 DEL PASS |
| 15 | 83550000 | <DEL> | PASS | END=83562000;SVTYPE=DEL;SVLEN=-12000 | 1/1 | chr15 83551620 83557670 6050 DEL PASS | |
| 15 | 84832500 | <DEL> | COMMON | END=84916500;SVTYPE=DEL;SVLEN=-84000 | ./. | chr15 84834155 84913077 78922 DEL NoConsensusGT | |
| 16 | 18828000 | <DEL> | PASS | END=18840000;SVTYPE=DEL;SVLEN=-12000 | 1/1 | | chr16 18832524 18838376 5852 DEL PASS |
| 19 | 21823500 | <DEL> | PASS | END=21835500;SVTYPE=DEL;SVLEN=-12000 | 0/1 | | |
| 19 | 35850000 | <DEL> | PASS | END=35863500;SVTYPE=DEL;SVLEN=-13500 | 0/1 | chr19 35849137 35861603 12466 DEL LongReadHomRef | |
| 19 | 43699500 | <DEL> | COMMON | END=43765500;SVTYPE=DEL;SVLEN=-66000 | ./. | chr19 43701473 43765338 63865 DEL NoConsensusGT | |
| 19 | 53517000 | <DEL> | PASS | END=53553000;SVTYPE=DEL;SVLEN=-36000 | 1/0 | chr19 53516912 53552060 35148 DEL NoConsensusGT | |
| 20 | 1560000 | <DEL> | COMMON | END=1587000;SVTYPE=DEL;SVLEN=-27000 | ./. | chr20 1552309 1585333 33024 DEL NoConsensusGT | |
| 20 | 54433500 | <DEL> | PASS | END=54445500;SVTYPE=DEL;SVLEN=-12000 | ./. | chr20 54434610 54440616 6006 DEL PASS | |
| X | 154789500 | <DEL> | PASS | END=154804500;SVTYPE=DEL;SVLEN=-15000 | ./. | chrX 154790189 154803271 13082 DEL ClusteredCalls | |
| Y | 22444500 | <DEL> | PASS | END=22458000;SVTYPE=DEL;SVLEN=-13500 | ./. | chrY 22446665 22458037 11372 DEL NoConsensusGT | |

### Supplementary Table 2. False positive deletions

| EventID | SvID | BreakID1 | BreakID2 | ChrA | PosA | ChrB | PosB | ShareBarcode | RealType | SimpleType | ComprehensiveFilter |
|---|---|---|---|---|---|---|---|---|---|---|---|
| S46 | 483 | 572 | 573 | 1 | 242482500 | 10 | 38562000 | 13 | LL | TRA3 | PASS |
| S46 | 484 | 572 | 574 | 1 | 242482500 | 10 | 38568000 | 13 | LR | TRA2 | PASS |
| S65 | 614 | 1860 | 818 | 2 | 133120500 | 9 | 68380500 | 32 | RL | TRA1 | PASS\|COMMON |
| S101 | 739 | 1031 | 3622 | 4 | 190662000 | 4 | 190683000 | 50 | RR | INV2 | PASS\|COMMON |
| S112 | 787 | 1188 | 3648 | 6 | 32460000 | 6 | 32557500 | 66 | RR | INV2 | PASS |
| S112 | 789 | 1192 | 3648 | 6 | 32526000 | 6 | 32557500 | 61 | RR | INV2 | PASS\|COMMON |
| S177 | 1132 | 2009 | 2010 | 10 | 81505500 | 10 | 81610500 | 121 | LR | DUP | PASS\|COMMON |
| S196 | 1160 | 2102 | 2103 | 12 | 17923500 | 12 | 18013500 | 109 | LL | INV1 | PASS\|COMMON |
| S196 | 1789 | 3760 | 3761 | 12 | 17922000 | 12 | 18010500 | 106 | RR | INV2 | PASS\|COMMON |
| S205 | 1176 | 2233 | 3769 | 14 | 19864500 | 14 | 20424000 | 93 | RR | INV2 | PASS\|COMMON |
| S205 | 1790 | 3769 | 3799 | 14 | 20424000 | 22 | 16131000 | 76 | RR | TRA4 | PASS\|COMMON |
| S224 | 1245 | 2537 | 3826 | 16 | 15123000 | 16 | 16408500 | 69 | RR | INV2 | PASS |
| S224 | 1795 | 3825 | 2537 | 16 | 15010500 | 16 | 15123000 | 112 | RR | INV2 | PASS\|COMMON |
| S226 | 1250 | 2600 | 2601 | 16 | 28432500 | 16 | 28705500 | 119 | LR | DUP | PASS\|COMMON |
| S255 | 1424 | 2931 | 2933 | 20 | 25827000 | 20 | 26070000 | 57 | LR | DUP | PASS\|COMMON |
| S288 | 1572 | 3300 | 3299 | X | 2698500 | Y | 2650500 | 81 | RL | TRA1 | PASS\|COMMON |
| S300 | 1796 | 3835 | 3836 | 16 | 21594000 | 16 | 22710000 | 187 | RR | INV2 | PASS\|COMMON |

**Supplementary Table 3. False positive inversions, duplications and translocation.**

| | | stLFRsv | stLFRsv | stLFRsv | stLFRsv+smoove | stLFRsv+smoove | stLFRsv+smoove |
|---|---|---|---|---|---|---|---|
| | Coverage | 30X | 50X | 100X | 30X | 50X | 100X |
| 50-1k | Benchmark | 4719 | | | | | |
| | Total-call | 0 | 0 | 0 | 359 | 649 | 972 |
| | True positive | 0 | 0 | 0 | 276 | 490 | 724 |
| | False positive | 0 | 0 | 0 | 83 | 159 | 248 |
| | False negative | 4719 | 4719 | 4719 | 4443 | 4229 | 3995 |
| | Precision | - | - | - | 76.88% | 75.50% | 74.49% |
| | Recall | - | - | - | 5.85% | 10.38% | 15.34% |
| 1k-10K | Benchmark | 577 | | | | | |
| | Total-call | 0 | 0 | 9 | 403 | 503 | 554 |
| | True positive | 0 | 0 | 9 | 325 | 405 | 434 |
| | False positive | 0 | 0 | 0 | 78 | 98 | 120 |
| | False negative | 577 | 577 | 568 | 252 | 172 | 143 |
| | Precision | - | - | 100.00% | 80.65% | 80.52% | 78.34% |
| | Recall | - | - | 1.56% | 56.33% | 70.19% | 75.22% |
| 10k-30k | Benchmark | 31 | | | | | |
| | Total-call | 16 | 26 | 60 | 16 | 26 | 60 |
| | True positive | 5 | 12 | 26 | 5 | 12 | 26 |
| | False positive | 11 | 14 | 34 | 11 | 14 | 34 |
| | False negative | 26 | 19 | 5 | 26 | 19 | 5 |
| | Precision | 31.25% | 46.15% | 43.33% | 31.25% | 46.15% | 43.33% |
| | Recall | 16.13% | 38.71% | 83.87% | 16.13% | 38.71% | 83.87% |
| >30k | Benchmark | 9 | | | | | |
| | Total-call | 13 | 16 | 18 | 13 | 16 | 18 |
| | True positive | 5 | 6 | 6 | 5 | 6 | 6 |
| | False positive | 8 | 10 | 12 | 8 | 10 | 12 |
| | False negative | 4 | 3 | 3 | 4 | 3 | 3 |
| | Precision | 38.46% | 37.50% | 33.33% | 38.46% | 37.50% | 33.33% |
| | Recall | 55.56% | 66.67% | 66.67% | 55.56% | 66.67% | 66.67% |

**Supplementary Table 4. Down sampled deletion evaluation on whole genome against GIAB HG002 benchmark**

| | Benchmark | Total-call | True positive | False positive | False negative | Precision | Recall |
|---|---|---|---|---|---|---|---|
| 50-1k | 6232 | 12410 | 4882 | 7528 | 1350 | 39.34% | 78.34% |
| 1k-10k | 909 | 1293 | 699 | 594 | 210 | 54.06% | 76.90% |
| 10k-30k | 25 | 7 | 5 | 2 | 20 | 71.43% | 20.00% |
| >30k | 6 | 0 | 0 | 0 | 6 | - | - |
| All | 7172 | 13710 | 5586 | 8124 | 1586 | 77.89% | 40.74% |

**Supplementary Table 5. Insertion evaluation for Nanopore long reads**
